# Targeting of the human nasal microbiota by secretory IgA antibodies

**DOI:** 10.1101/2022.08.31.505993

**Authors:** Rob van Dalen, Ahmed M. A. Elsherbini, Mareike Harms, Svenja Alber, Regine Stemmler, Andreas Peschel

**Affiliations:** Interfaculty Institute of Microbiology and Infection Medicine, Infection Biology, University of Tübingen, Tübingen, Germany

**Keywords:** mucosal immunology, secretory IgA, IgA-seq, metagenomics, nasal microbiome

## Abstract

The human nasal microbiome is critical for health and disease, since it is associated with the occurrence of respiratory disorders and hosting of opportunistic pathogens. The host therefore protects this vulnerable mucosal barrier from infection and maintains homeostasis of the microbiota through various mechanisms, including the production of secretory IgA (sIgA) antibodies. However, we currently lack a comprehensive understanding of how sIgA affects the nasal microbiota. Through IgA-seq analysis of nasal microbiome samples and sIgA deposition experiments using nasal sIgA from healthy volunteers, we identified which bacterial genera and species are targeted by sIgA on the level of the individual host. We observed that the amount of sIgA secreted into the nasal mucosa by the host varied substantially and was negatively correlated with the bacterial density. The interaction between mucosal sIgA antibodies and the nasal microbiome was highly individual, and was not dependent on the microbiome composition, or the age or gender of the host. Importantly, we showed that for the clinically relevant opportunistic pathogen *S. aureus*, sIgA reactivity was in part the result of epitope-independent interaction of sIgA with the antibody binding protein SpA through binding of sIgA Fab regions. This study thereby offers a first comprehensive insight of targeting of nasal microbiota by sIgA antibodies, which may help to better understand the shaping and homeostasis of the nasal microbiome by the host and offer new targets for intervention in disease-associated microbiota.

## Introduction

The human nasal microbiome is an important factor in health and disease. Its composition is associated with respiratory disorders, such as chronic rhinosinusitis and allergies, and it can additionally host opportunistic pathogens such as *Staphylococus aureus* (1, 2). Compared to the gut microbiome, the nasal microbial community is relatively scarce. In most individuals, it comprises of only two to ten bacterial species (3). The community composition varies strongly between individuals, but can be divided into seven general profiles. These community state types (CSTs) are defined based on the presence of several hallmark members of the community, like *S. aureus, Staphylococcus epidermidis, Cutibacterium* spp. or *Corynebacterium* spp. (4). Due to the presence of opportunistic pathogens in the nose, the host needs to protect this vulnerable mucosal barrier from infection and maintain homeostasis of the local microbiota. It can achieve this by several means, including limitation of nutrient availability in the nasal cavity in a process known as nutritional immunity (5), and through the production of antimicrobial defense proteins and peptides such as lysozyme, lactoferrin and α- and β-defensins (6, 7). Another hallmark feature of mucosal tissues, including the nasal mucosa, is the production of secretory IgA (sIgA) antibodies (8).

sIgA is a dimeric antibody that is highly abundant in nasal secretion, saliva, sweat, gut fluid, tears and milk, and is distinctively different from the monomeric IgA found in human serum (8). sIgA contributes to the defense of the mucosa through various mechanisms. For instance, sIgA can prevent interaction of pathogens with the epithelium by agglutinating them and blocking their adhesion molecules, in a process known as immune exclusion (8). This was observed in the gastrointestinal tract, in which sIgA coated and facilitated clearance of gut bacteria that are associated with the onset of colitis, thereby providing protection against disease (9). Paradoxically, sIgA can also facilitate bacterial colonization of the mucosa, by enhancing the mucus-binding properties of the bacterial cell surface through sIgA coating, such as is the case for the prominent human gut commensal *Bacteroides fragilis* (10, 11). These contrasting effects of sIgA coating of bacteria are mediated by the mucus flow rate, with low mucus flow rates facilitating colonization and high rates facilitating clearance (11). These important insights into the role of sIgA in the control of the microbiota were gained particularly through studies on sIgA in the gastrointestinal tract. Especially the development of the IgA-seq technology, in which microbiota are sorted based on being coated by sIgA or not and subsequently analyzed by metagenomic sequencing (9, 12, 13), has significantly contributed to our understanding of the role of sIgA in the homeostasis of the microbiome and defense against pathogens in the gastrointestinal tract (9, 10, 12, 14).

However, we currently still lack a comprehensive understanding of how sIgA affects microbiota in other mucosal niches, in particular in the human nose. A role of sIgA in the control of the nasal microbiome is implied by the observation that individuals with selective IgA deficiency frequently suffer from allergies and recurrent respiratory infections (15, 16). Moreover, various respiratory pathogens produce immune evasion factors that inhibit sIgA, including an IgA serine protease produced by *Haemophilus influenzae*, the IgA-binding protein SSL7 secreted by *S. aureus*, and the secreted antibody lambda-chain binding protein Protein L by *Finegoldia magna* (17–21). This suggests a benefit for these species to evade sIgA immune responses and thereby a role for sIgA in the defense against these pathogens.

*S. aureus* is of particular interest in this regard, as it is part of the normal human nasal microbiome, colonizing approximately one-third of the human population permanently and one-third intermittently (22). However, nasal colonization by *S. aureus* is also an important risk factor for life-threatening infections (23). Evasion of antibody responses is an important virulence strategy of *S. aureus*. Other than the secreted SSL7, it produces the surface proteins staphylococcal protein A (SpA) and the second staphylococcal immunoglobulin-binding protein (Sbi) factors that bind various classes of antibodies. SpA is well-characterized to bind the Fc region of most human antibody classes, although not Fc of IgA. However, through different binding sites SpA also binds Fab domains of antibodies that belong to the structural VH3 family (24, 25). Although Fab-mediated binding by SpA has to our knowledge not been reported for IgA or sIgA specifically, the structure of the sIgA molecule should not preclude this interaction from taking place. Sbi, on the other hand, does not interact with sIgA as it only binds the Fc region of the IgG antibody class (26, 27). Although the role of *S. aureus* antibody binding proteins in invasive disease is well studied, their role in the context of nasal colonization is currently still unknown.

In this study, we aimed to determine whether sIgA affects the nasal microbiome composition. We therefore applied IgA-seq on nasal microbiome samples from healthy adult volunteers to identify which bacterial species are targeted by sIgA, with a particular interest for *S. aureus*.

## Results

### Study population and nasal community state types

Before analyzing the sIgA-binding capacities of nasal bacteria, we determined the overall nasal microbiome composition of the study participants in a cohort of 50 healthy human volunteers (**Table 1**), using a previously published method for 16S rRNA sequencing that has been optimized for the human nasal microbiome (3, 28). All study participants were classified into the nasal CSTs (**Figures 1A, B**), using a custom Python classifier script based on the original CST definitions by Liu *et al*. (4). Strikingly, we observed a much higher proportion of the *Corynebacterium* spp.-defined CST5 in our study population compared to Liu *et al*. (54.0% vs. 20.0%) (4). In contrast, we observed a lower prevalence of the *S. aureus*-defined CST1 (4.0% vs. 12.4%), *S. epidermidis-defined* CST3 (16.0% vs. 22.5%) and *Cutibacterium* spp.-defined CST4 (16.0% vs. 28.7%). Additionally, we found no Enterobacteriaceae-defined CST2, *Moraxella* spp.-defined CST6 and *Dolosigranulum pigrum*-defined CST7 in our study population at all. Five samples (10%) could not be classified into any of the CSTs and were therefore labeled as ‘unclassified’. Consistent with the work of Liu *et al*. (4), we detected no differences in the microbiome composition between the genders (**Figure S1**). Of note, the alpha diversity of the highly prevalent CST5 was higher compared to that of CST4 (**Figure 1C**), indicating that *Corynebacterium* spp. supports a broader microbial composition than *Cutibacterium* spp. In line with the findings of Escapa *et al*. (3), *Corynebacterium, Cutibacterium* and *Staphylococcus* were the most prevalent and abundant genera in our study population. In contrast, we detected *Lawsonella clevelandensis* only in a single individual, despite it being described previously as a highly prevalent nasal species (3).

**Figure 1.**
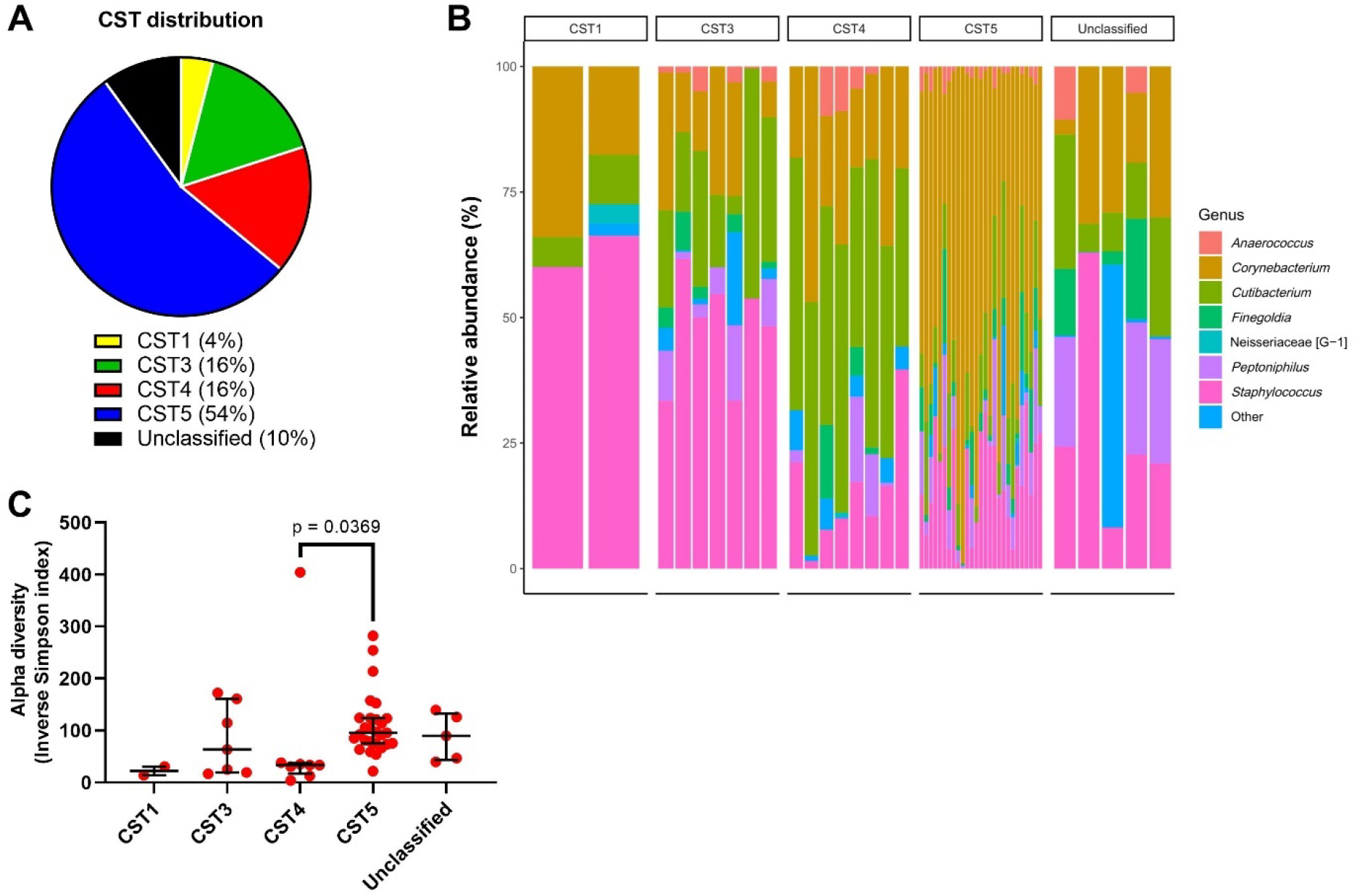
CST classification of the study population. (**A**) Study participants were classified into the previously described CSTs, based on the non-sorted microbiome samples. Samples that did not fit into any of the seven originally defined CSTs were labeled as ‘unclassified’. (**B**) Relative abundances of the most common species and genera as well as the CST indicator species and genera in all study participants, stratified by CST. (**C**) Alpha diversity expressed as the Inverse Simpson index of all identified CSTs. Statistical differences in the alpha diversity were calculated by Kruskal-Wallis test with Dunn’s test for multiple comparisons.

**Table 1.**
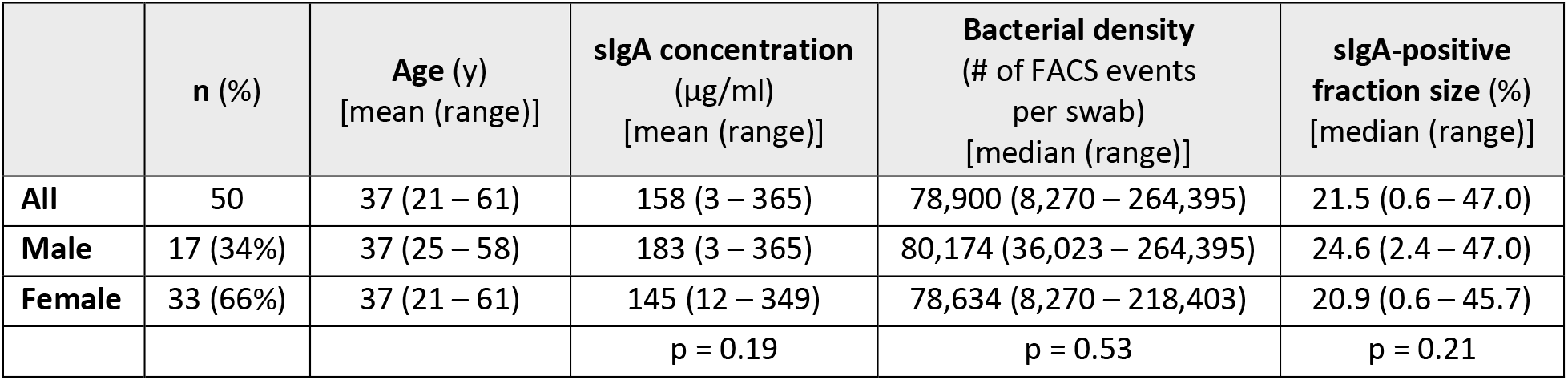
Study population and FACS sorting characteristics. Statistical differences between the two genders were tested by Mann-Whitney test (for bacterial density and fraction size) or unpaired t-test (for sIgA concentration).

The unclassified samples contained large proportions of a yet-unnamed *Peptoniphilus* species designated as human microbial taxon (HMT) 187, *Streptococcus* HMT-064 or *Staphylococcus capitis* (**Figures 1B, S2A**), suggesting the possibility of yet undefined CSTs. Principal coordinates analysis of the beta diversity revealed that two of the unclassified samples (including the sample with a high proportion of *S. capitis)* cluster with the *S. aureus*-defined CST1 samples, whereas the other three unclassified samples cluster with CST3 and CST5 (**Figures S2B, S2C**). This indicates that these community compositions could be variations of these CSTs that are not included in the current CST definitions.

### Nasal sIgA limits the nasal bacterial density

We next determined the sIgA concentration of nasal swab eluates from all study participants. This showed a large variation in sIgA quantity between individuals, ranging from 3 to 365 μg/ml (**Figure 2B**). Since the swabs were eluted in a volume of 1 ml, these concentrations can be interpreted as the absolute quantities of sIgA collected using a single swab. To elucidate how human nasal sIgA may target the full range of nasal bacteria and contribute to microbiome diversity, bacteria eluted from human nasal swabs were stained with a fluorescently-labeled antibody specific for human IgA, thereby labeling pre-deposited sIgA on native microbiome samples. The bacteria were subsequently sorted by FACS according to their fluorescence into sIgA-positive and sIgA-negative fractions (**Figure 2A**). Using the total FACS event count per swab as a measure for the absolute bacterial density (**Figure 2C**), we observed that the nasal sIgA concentration correlates negatively with the bacterial density (**Figure 2E**). In contrast, the nasal sIgA concentration shows a positive correlation with the proportion of FACS events sorted into the sIgA-positive fraction (**Figures 2D, 2F**). These observations suggest that, when considering the nasal microbiome as a whole, sIgA coating of bacteria negatively affects the bacterial density. However, this does not consider any differences in sIgA coating between the bacterial species present in the microbiome or the subsequent functional effects of this (either supporting bacterial clearance or colonization). Furthermore, we found no correlation between the sIgA concentration and the alpha diversity, indicating that sIgA did not impact the diversity of the nasal microbiome (**Figure 2G**). Of note, no differences in the bacterial density, sIgA-positive fraction size or sIgA concentration between males and females were identified (**Table 1**), in contrast to a previous report of a higher nasal bacterial density in males (4). Taken together, this indicates that sIgA limits the overall nasal bacterial density, but not the diversity.

**Figure 2.**
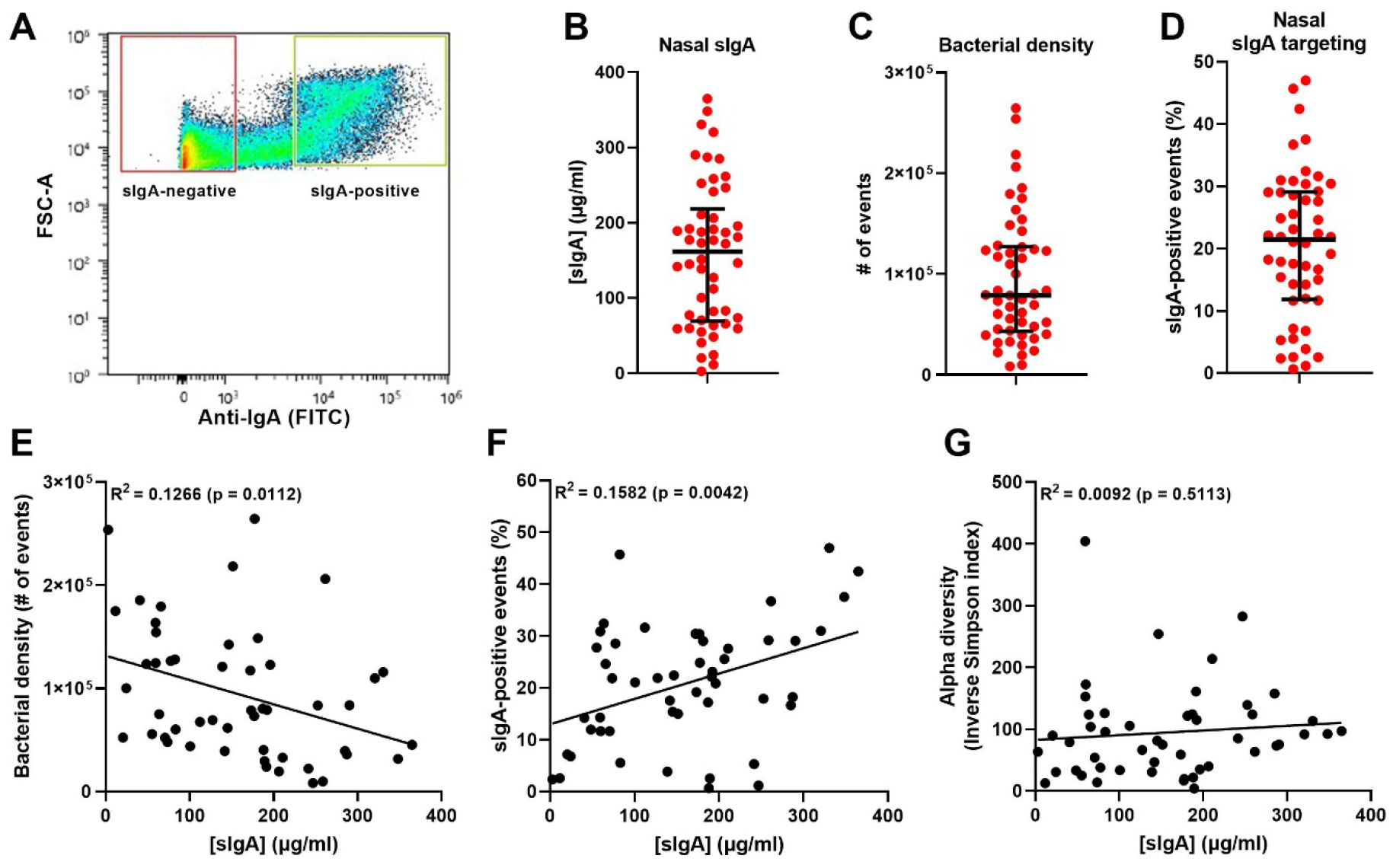
Nasal sIgA concentration and correlation with bacterial density and diversity. (**A**) Sorting by FACS of a representative nasal microbial sample into sIgA-positive (green gate) and sIgA-negative (red gate) fractions, based on forward scatter (FSC-A) and anti-IgA staining. (**B-D**) Distribution with the median ± interquartile range of (**B**) nasal sIgA concentrations in the filtered nasal eluates of all study participants, (**C**) nasal bacterial density, expressed as the total number of events analyzed by FACS, and (**D**) percentage of sIgA-positive events detected by FACS. (**E-G**) Correlation of the nasal sIgA concentration with (**E**) the bacterial density expressed as the total number of FACS events, (**F**) the percentage of sIgA-positive events detected by FACS, and (**G**) the alpha diversity expressed as the Inverse Simpson index in the non-sorted fraction. Correlations were tested by linear regression analysis.

### Highly individual sIgA-targeting of nasal microbiome members

To analyze if nasal sIgA targets all nasal bacteria equally effectively or tends to target specific species in a given host, the microbial composition of the sIgA-positive and sIgA-negative samples were determined as before. After exclusion of three samples (one each of CST1, CST3 and CST5) for which 16S rRNA sequencing failed due to too limited DNA yields, we observed that bacterial sorting based on sIgA coating resulted in a substantially altered sample composition. The alpha diversity of the sIgA-positive and sIgA-negative fractions was significantly reduced compared to the non-sorted samples (**Figure 3A**). IgA probability ratios (‘IgA scores’ from here on) were calculated for all genus-level and species-level taxa (13) (**Supplementary file**). These scores are a measure for sIgA targeting of specific taxa within an individual, and range from −1 (all bacteria of a given taxon are in the sIgA-negative fraction) to +1 (all bacteria of a given taxon are in the sIgA-positive fraction). We selected the top-20 species and the top-6 genera with the highest frequency of IgA scores for further analysis (**Figure S3**). For the majority of genus-level or species-level taxa, we observed a highly variable IgA score that in several taxa varied across almost the entire range from −1 to +1 among the study population (**Figures 3B, 3C**). This indicates highly variable sIgA responses to the microbiota by the individual hosts. On genus-level, there was an overall trend towards negative IgA scores, with *Staphylococcus, Cutibacterium* and *Corynebacterium* having IgA scores significantly below 0, indicating that these genera were generally not effectively targeted by nasal sIgA antibodies (**Figure 3B**). On species-level, we again observed an overall trend towards negative IgA scores, although only two species (*Cutibacterium acnes* and *Paracoccus yeei*) had IgA scores significantly below 0 (**Figure 3C**). Interestingly, for the majority of species the IgA scores were not normally distributed across the IgA score range of −1 to +1 (17 out of 20 species with a Shapiro-Wilk test outcome of p < 0.05), but instead formrf three distinct clusters of negative, zero-centered or positive IgA scores (**Figure 3C**). This is the case for e.g. *S. aureus, S. epidermidis* and *Corynebacterium accolens*. Since every datapoint represents an individual host, this indicates three different main modes of sIgA interactivity with these species across individuals, in which only several individuals produce an effective sIgA response against particular microbiota, whereas others do not effectively target these microbiota at all. Strikingly, other species, such as *C. acnes*, lack this separation into three clear clusters and show a more closely-clustered distribution of IgA scores (**Figure 3C**).

**Figure 3.**
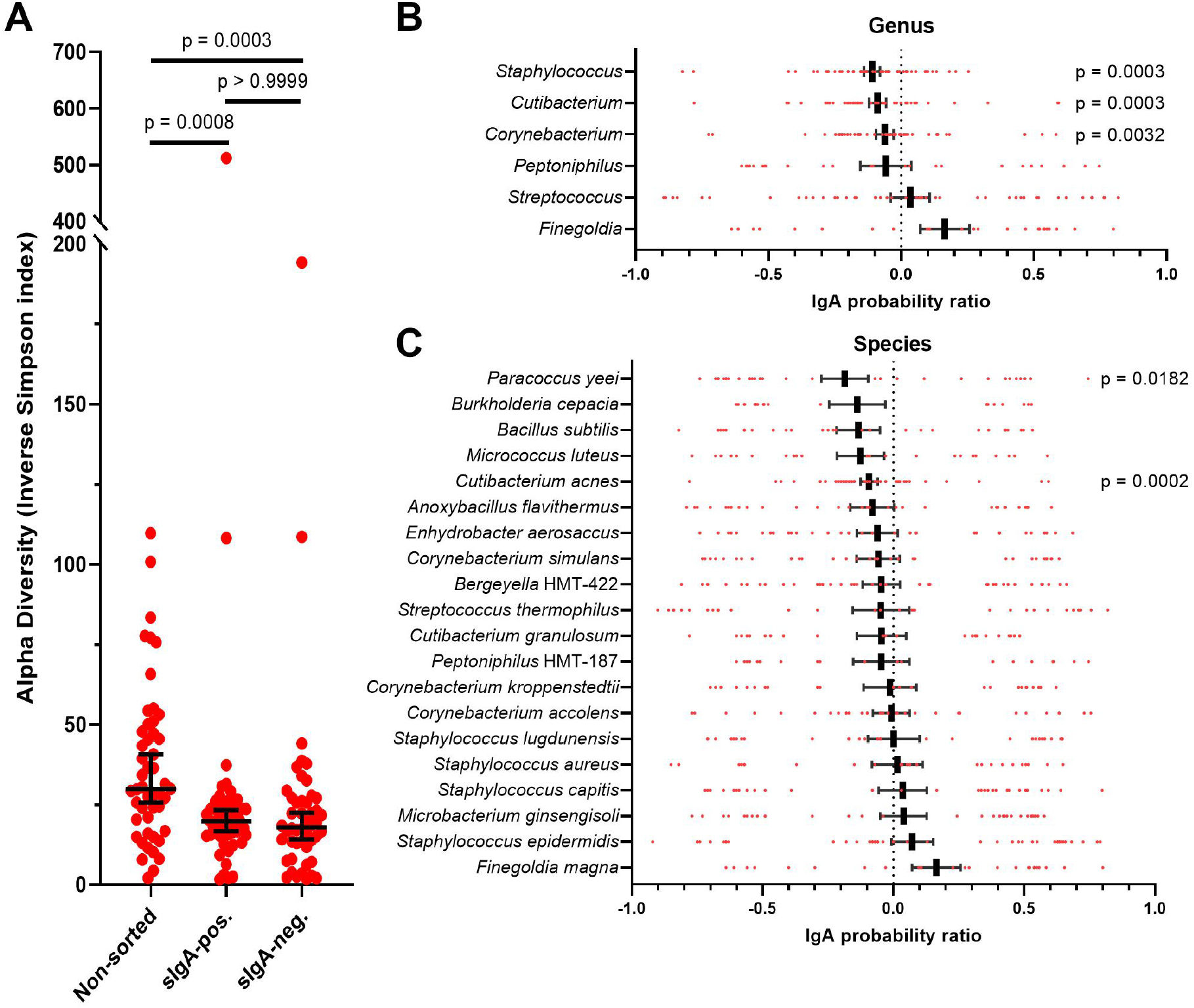
IgA targeting of nasal bacteria. (**A**) Alpha diversity of the non-sorted, sIgA-positive and sIgA-negative fractions, expressed as the Inverse Simpson index. Individual values are depicted in red, with the median and 95% confidence interval in black. Statistical differences were calculated by Kruskal-Wallis test with Dunn’s test for multiple comparisons. (**B-C**) sIgA binding is presented as the IgA probability ratio on a scale from −1 to +1 for (**B**) the six most prevalent genera and (**C**) the 20 most prevalent species (also see **Figure S3**). Individual values are depicted in red, with the mean ± SEM in black. IgA scores were tested by Wilcoxon signed-rank test against a hypothetical value of 0.

We next examined how sIgA targeting varied on the individual host level, by hierarchical cluster analysis of the IgA scores for both the hosts and the previously-selected top-20 species (**Figures 4, S4**). Based on this analysis, several clusters of hosts with distinct sIgA profiles could be distinguished. Branches 1 and 3 contained individuals who produce sIgA that broadly covers their nasal microbiome, as indicated by the generally positive IgA scores of these individuals. In contrast, sIgA of the individuals in branch 2 was generally poorly reactive with the nasal microbiota of these individuals, as indicated by the generally negative IgA scores. Branches 4 and 5 instead contained individuals with varying levels of sIgA reactivity to different species present. Importantly, sIgA targeting did not depend on the *S. aureus* carrier status, nasal sIgA concentration, age, gender or CST of the host, since no clustering of these factors with the IgA scores was observed (**Figure 4**). These results suggest a highly individualized interplay between our immune system and our nasal microbiota that is irrespective of the resident nasal microbiome.

**Figure 4.**
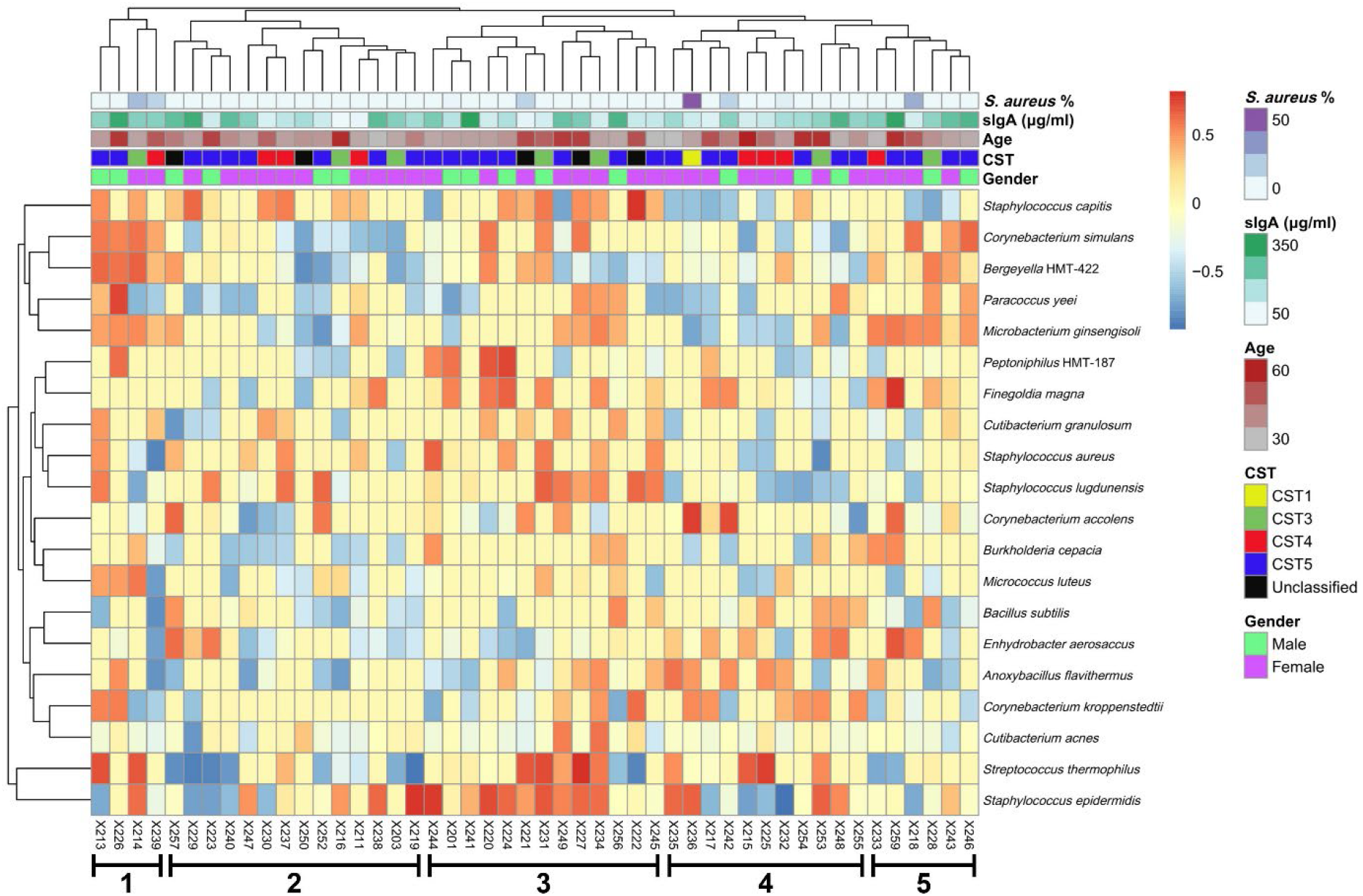
Hierarchical cluster analysis of the sIgA targeting. The study participants (columns) and the top-20 nasal bacterial species (rows) were clustered based on the IgA scores. Branches that contain hosts with distinct sIgA repertoires are indicated and numbered. See **Figure S4** for the same analysis expanded to all detected nasal bacterial species.

### Validation of sIgA-targeted nasal species

To validate the IgA-seq approach, we measured by flow cytometry sIgA deposition on *in vitro* cultures of a selection of representative strain isolates from nasal bacterial species that were prevalent in this study (**Figures 5A, B**). As sIgA source we used individual sterile-filtered nasal swab eluates collected from the study participants and that were adjusted to the same sIgA concentration. For each species analyzed this way, the signal of the deposited sIgA on the bacteria was subsequently plotted against the IgA score of the same species from the corresponding study participant. Using this method, we could validate the nasal IgA-seq approach, as we observed a correlation between the sIgA deposition and the IgA score for *S. epidermidis, Staphylococcus lugdunensis, C. accolens* and *C. acnes* (**Figure 5A**). In all cases, the individuals with the highest IgA scores also ranked among the highest for sIgA deposition. However, for all species we also observed a loss of resolution in the sIgA deposition for individuals with IgA scores below 0.5. This is indicative of a lower sensitivity of the sIgA deposition assay, particularly in the case of poorly-reactive sIgA preparations. Regardless, for these species we could broadly validate the IgA-seq outcomes.

**Figure 5.**
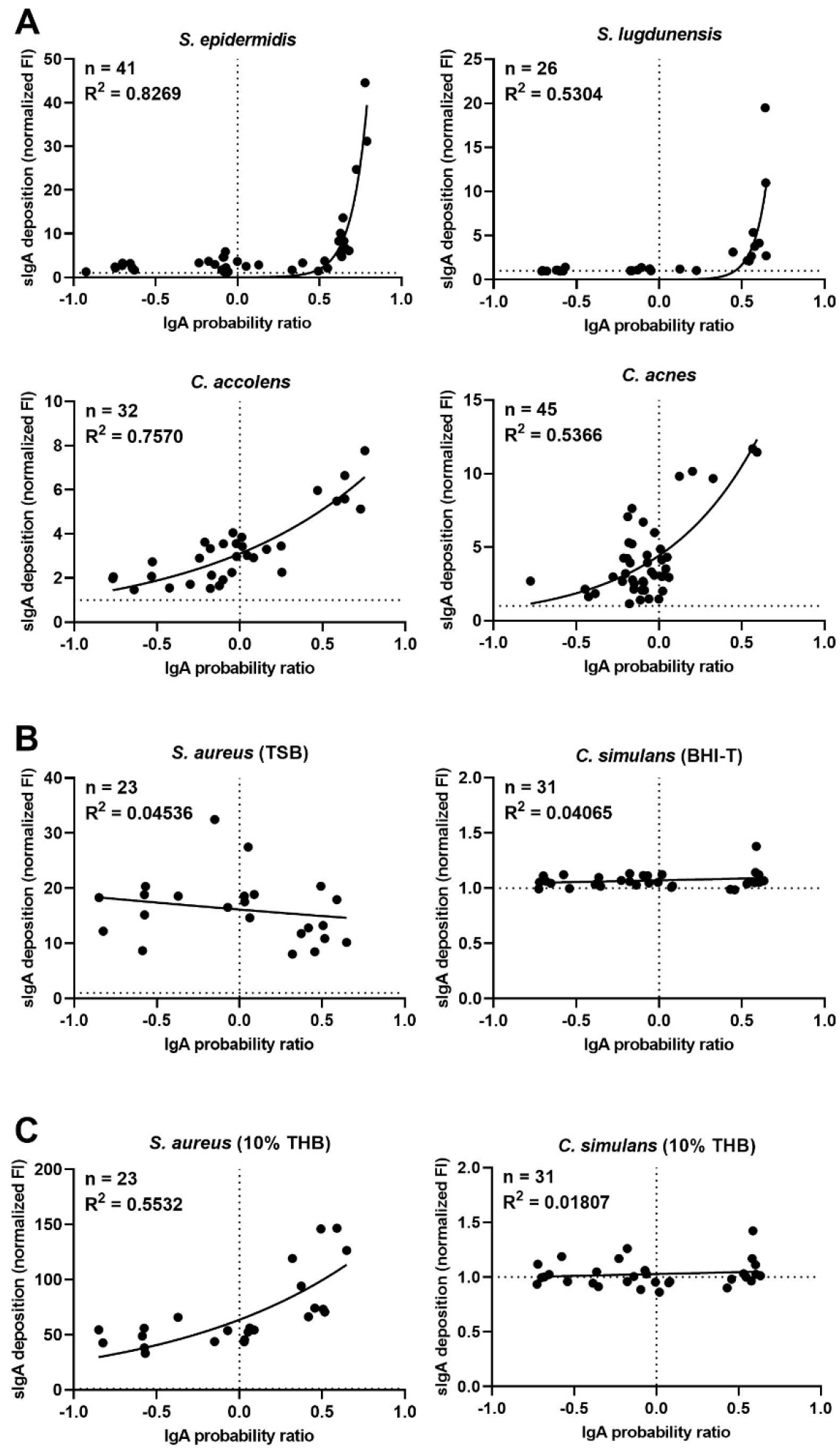
Validation of the IgA-seq outcomes. (**A, B**) Correlation between sIgA deposition from filtered and concentration-adjusted nasal eluates on cultured nasal bacterial species, as assessed by flow cytometry, and the IgA probability ratio calculated for the same species of the matching study participants. All strains were grown in nutrient-rich TSB (for *Staphylococcus)* or BHI-T (for *Corynebacterium* and *Cutibacterium)*. All samples were normalized to 10 μg/ml sIgA or 0.3 μg/ml sIgA for *S. aureus*. Fluorescence intensity (FI) was measured by flow cytometry in triplicates and normalized to the corresponding unstained controls, with a fold change of one (no detectable deposition) being represented by a dashed horizontal line. Correlations were analyzed by non-linear regression. (**C**) Correlation between sIgA deposition and IgA probability ratio for *S. aureus* and *C. simulans* grown in nutrient-limiting 10% THB.

Two major exceptions from these observations were *S. aureus* and *Corynebacterium simulans*, for which no correlation between the IgA score and the sIgA deposition was found (**Figure 5B**). In case of *S. aureus*, the universally high levels of *S. aureus*-reactive sIgA antibodies in the nasal swab eluates required the use of 30-fold lower sIgA concentration in the assay, compared to all other tested species. Despite being measured at a lower sIgA concentration, the sIgA deposition was much higher on *S. aureus* compared to any other species. On the other hand, for *C. simulans* we could hardly detect any sIgA deposition at all. For both *S. aureus* and *C. simulans*, this is potentially the result of large differences between the epitope repertoires produced by the representative strain isolates used in the sIgA deposition assay and the native strains colonizing the nares of the study participants. Alternatively, the nutrient-rich broths used for the *in vitro* cultivation of the isolates used in the sIgA deposition assay could potentially induce different epitope repertoires compared to the nutrient-poor conditions encountered in the human nares. This would result in sIgA deposition profiles that are not representative of the natural niche and therefore be poorly comparable to the IgA scores of native samples. To test this hypothesis, we grew the same *S. aureus* and *C. simulans* isolates in nutrient-limited 10% Todd Hewitt broth (THB) (29). Indeed, in stark contrast with *S. aureus* grown in nutrient-rich TSB, sIgA deposition on *S. aureus* grown in nutrient-limited 10% THB correlated with the IgA scores obtained by IgA-seq (**Figure 5C**). This effect was specific for *S. aureus*, as sIgA deposition on *C. simulans* was still virtually absent. This suggests that the nasal sIgA antibodies specific for *S. aureus* are at least partially dependent on epitopes that strongly vary depending on the growth condition.

### *S. aureus* binds sIgA aspecifically through SpA

For the individuals with low *S. aureus* IgA scores, we observed substantial sIgA deposition on *S. aureus* (**Figures 5B, 5C**). Since these observations seem to contradict each other, we decided to analyze sIgA deposition on *S. aureus* in the study participants for whom no *S. aureus* IgA score could be calculated due to the absence of this species in their microbiome. To our surprise, deposition of nasal sIgA from these individuals on *S. aureus* was at a similar level as that of the study participants who were colonized with *S. aureus* (**Figure 6A**). This suggests that nasal sIgA reactive with *S. aureus* is a universal phenomenon.

**Figure 6.**
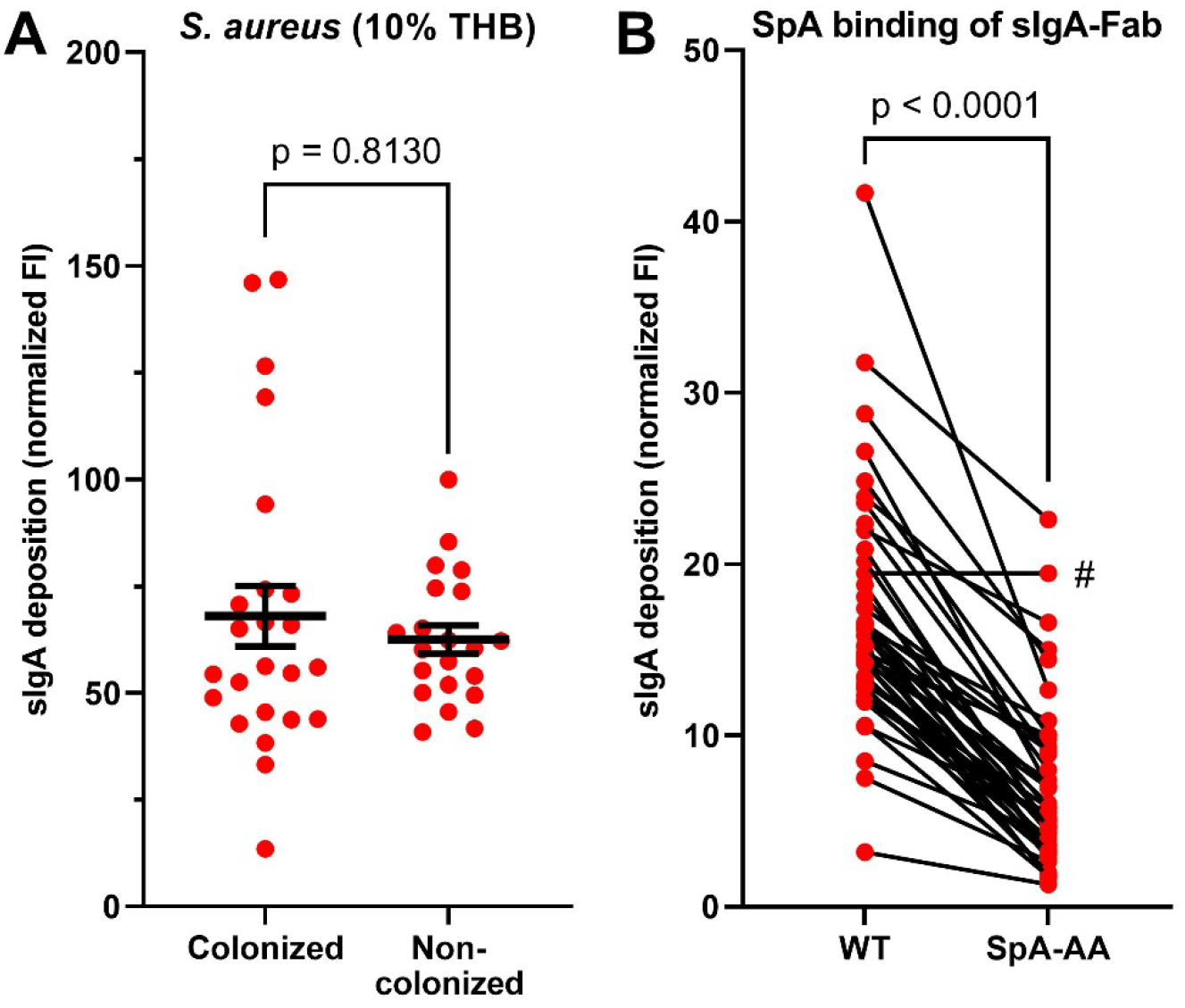
*S. aureus* binds sIgA aspecifically via SpA. (**A**) sIgA deposition of nasal sIgA of individual study participants on a nasal *S. aureus* isolate grown in 10% THB, stratified by the presence or absence of *S. aureus*. Statistical differences were tested by Mann-Whitney test. (**B**) sIgA deposition of nasal sIgA of individual study participants on *S. aureus* USA300 JE2 WT and the SpA-AA mutant deficient for binding of antibody Fab regions. Statistical differences were tested by paired t-test. A single sample (#) did not follow the trend of lower sIgA deposition on the SpA-AA mutant.

Since *S. aureus* is well-known to produce multiple antibody-binding proteins such as SpA, Sbi and SSL7 (18, 25, 27), we hypothesized that the observed universal sIgA deposition on *S. aureus* was not be epitope-based, but rather a result of epitope-independent binding of sIgA. SpA was a likely candidate to be this interaction partner, because, although unable to bind the Fc region of human IgA, it has the potential to bind the Fab region of antibodies that belong to the VH3 structural family (24). Using an *S. aureus* SpA-AA mutant that is deficient for VH3 Fab binding (25), we indeed observed a significant reduction in deposition of sIgA (**Figure 6B**). This reduction could be observed for nasal sIgA from all but one of the study participants, indicating the presence of VH3 sIgA antibodies on the nasal mucosa. The expression of SpA by *S. aureus* in the human nares is known to be highly variable between individuals and often distinctly different from conditions used *in vitro* (30). We therefore speculate that for the individuals with negative IgA scores but with substantial sIgA deposition *in vitro*, the colonizing *S. aureus* strains produce SpA at only low levels in the nares. Overall, these data indicate that deposition of nasal sIgA on *S. aureus* was largely dependent on epitope-independent SpA-sIgA interactions.

## Discussion

We currently lack a comprehensive understanding of how sIgA affects microbiota and supports microbiome homeostasis and immune defense in the human nasal mucosa. To determine whether sIgA affects the nasal microbiome composition, we applied IgA-seq on nasal microbiome samples from healthy adults to identify which bacterial species are targeted by sIgA. We observed highly individualized interplay between mucosal sIgA antibodies and the nasal microbiome. Importantly, we showed that for the highly prevalent and clinically relevant opportunistic pathogen *S. aureus*, sIgA reactivity was in part dependent on epitope-independent interaction with the antibody binding protein SpA.

On the host side, humans differed substantially (up to 100-fold) in the quantity of nasal sIgA that is produced during healthy conditions. Similar ranges of nasal sIgA concentrations have been reported previously by others (31–33). When considering the nasal bacterial community as a whole, high nasal sIgA amounts limited nasal bacterial density but not the microbial diversity. Importantly, high sIgA amounts did not favor particular CSTs or specific species. On the level of the bacterial species, IgA-seq revealed highly varying individual nasal sIgA profiles that were not dependent on age, sex or CST. Strikingly, for many species such as *S. aureus, C. accolens* and *Finegoldia magna*, individuals fall into one of three distinct IgA score groups of negative, zero-centered or positive IgA scores. This hints at three main modes of sIgA interactivity with these species, in which only several individuals produce a clear sIgA response against particular microbiota, whereas others do not target these microbiota at all or only mildly. Importantly, through cluster analysis we identify clusters of hosts producing an sIgA repertoire that broadly covers their nasal microbiota, whereas others are generally poorly reactive to their nasal microbiota or display varying levels of sIgA reactivity to different species of their microbiome. This suggests that sIgA responses generally depend on the host’s immune system rather than the immunogenicity of the colonizing bacterial species, implying a highly individualized interplay between the immune system and our microbiota. However, this complicates defining a single function of sIgA in the nasal cavity. In the gastrointestinal environment, sIgA can have both pro-microbial and anti-microbial properties, depending on mucus flow rate, disease state and differential adhesion between species (9–11). For the nasal niche, an important next step in this regard would be to compare baseline sIgA targeting with that of diseased states, such as chronic rhinosinusitis, cystic fibrosis, antibiotic-induced dysbiosis or other respiratory disorders. Furthermore, functional studies are required to define how sIgA coating affects bacteria in the nasal mucosa, taking into consideration the unique properties of this particular niche, such as the mucus flow rate and the exposure to the environment.

On the bacterial side, we did not detect any species that is universally targeted by sIgA. Both invasive and non-invasive bacteria show reactivity with sIgA without a notable pattern across the tested hosts. Therefore, bacterial aggressiveness is seemingly not correlated to the induction of nasal antibody responses. Particularly in the context of nasal vaccines, this is an important notion as it could preclude the use of bacterial vaccination platforms. A potential exception to this is *F. magna*, which showed the highest sIgA reactivity of any species or genus, though not significant in our study population. Although *F. magna* typically does not cause infection in healthy individuals, it is an opportunistic pathogen that can cause significant morbidity in immunocompromised hosts or hosts with barrier disruptions, ranging from skin abscesses to bone and prosthetic joint infections (34, 35). Additionally, it has the potential to activate neutrophils and trigger NETosis (36). These features could explain the high sIgA reactivity to *F. magna*. Alternatively, the observed sIgA targeting could also be partially or fully mediated by the *F. magna* antibody-binding Protein L and therefore be epitope-independent (20, 21), although this requires further investigation.

In the case of *S. aureus*, we did confirm epitope-independent interaction with sIgA to take place. *S. aureus* showed high deposition of sIgA from all study participants, despite not being universally sIgA-targeted in the IgA-seq approach. We determined that this high level of interaction was caused for a significant part by epitope-independent interaction of antibody-binding protein SpA with sIgA. Since SpA cannot bind IgA Fc, this interaction was dependent on Fab-binding of VH3-family sIgA (25). We excluded the involvement of other antibody-binding proteins of *S. aureus* (Sbi and SSL7) based on their inability to bind IgA or for not being cell-surface bound, respectively (18, 26, 27). SpA involvement in this phenomenon was further supported by the increase in sIgA deposition of *S. aureus* in nutrient-limited growth conditions, under which there is a lack of *spa* repression by the *agr* quorum sensing system (30) and thus an increase in sIgA binding. Importantly, *spa* expression in the human nares is highly variable between individuals and significantly different from *spa* expression by the same strains *in vitro* (30). This provides a rational for the negative *S. aureus* IgA scores we observed in a number of individuals, despite the sIgA of all individuals showing the potential to deposit on the *S. aureus* cell surface under laboratory conditions. However, further studies are required to confirm this. Importantly, the functional consequences of SpA-mediated sIgA coating of *S. aureus* in the human nasal mucosa is currently unknown. Based on the function of sIgA in the gastrointestinal tract (11), these could range from improved *S. aureus* clearance from the nasal cavity, to improved colonization through enhanced mucus- or epithelial interactions. Intentionally coating itself in host immune factors to gain benefit would not be uncommon for *S. aureus*, which is known to coat itself in fibrinogen through Efb (37). Further research on the functional consequences of SpA-sIgA interaction on the human nasal mucosa is therefore necessary.

We also observed several striking differences between our data and the previously defined nasal CSTs. The classification of microbial community compositions into CSTs is an efficient method to reduce dimensionality of the data and it provides an easy label to describe a particular microbial community. To improve reproducibility of nasal CST classification, we used a custom script that takes an easy to use but more reductive approach (only the most abundant species or genus is considered, rather than the whole community) to CST classification compared to the original publication (4). The nasal microbiomes of our study population were generally in line with previously published data (3, 4), but we observed several striking differences in CST prevalences. In particular, none of the analyzed microbiomes classified as the *M. catarrhalis-characterized* CST6, *D. pigrum-characterized* CST7 or Enterobacteriaceae-characterized CST2, as these taxa were relatively rare in our population (4). In contrast, our study population has a large proportion of the *Corynebacterium*-characterized CST5 (54% of individuals). Interestingly, multiple microbiomes (10%) with a high abundance of *Peptoniphilus* HMT-187, *Streptococcus* HMT-064 or *S. capitis* could not be classified into any of the CSTs. Beta diversity analysis suggested that these microbiomes could be variations of other CSTs, expanding their current definitions. Alternatively, it is possible that these microbiome compositions are actually representative of currently unrecognized CSTs, although further studies will be necessary to confirm this.

Lastly, our results stress the importance of laboratory testing of bacteria in conditions that mimic their niche environment. The use of nutrient-rich culture media in general research practice will often hide phenomena that depend on the unique conditions in the body niches that these bacteria grown in, broadly affecting regulatory mechanisms, metabolic adaptation and expression of epitopes (5, 30, 38). Measuring microbial processes or host-microbe interaction in the native state in their niche, such as by IgA-seq, is therefore essential to better understand colonization, infection and microbiome homeostasis. IgA-seq has additionally hinted at the potential significance of low-abundant nasal species, such as *Paracoccus yeei*.

In conclusion, the nasal IgA-seq approach revealed highly individualized interplay between mucosal sIgA antibodies and the nasal microbiome. Thereby, this study offers a first comprehensive insight of targeting of nasal microbiota by sIgA antibodies, which can aid in better understanding of the shaping and homeostasis of a healthy nasal microbiome by the host. In the case of the important pathogen *S. aureus*, sIgA reactivity was epitope-independent due to binding of sIgA by the antibody binding protein SpA. Together, these interactions may offer new targets for intervention in disease-associated microbiota.

## Methods

### Human sample collection and processing

This study was approved by the Institutional Review Board for Human Subjects at the University of Tübingen (143/2020BO2) and informed written consent was obtained from all healthy adult volunteers before sample collection.

From each study participant we sampled both anterior nares using E-Swabs (Copan Diagnostics) that were briefly dipped into sterile PBS (Lonza), and were stored immediately in 1 ml of the provided Amies transport medium at 4°C for up to 18 hours, according to the manufacturer’s instructions. To ensure sample-to-sample reproducibility, all samples were collected by the same researcher following the same procedure for each sample: swabs were swirled 10 times around the outer rim of each nostril at a depth of 1-2 cm.

Bacteria were eluted from the swab, centrifuged (1 minute, 10 000 xg), aspirated and resuspended in 1 ml PBS. Per sample, a 50 μl aliquot was taken as unstained control for fluorescence-activated cell sorting (FACS) and a 200 μl aliquot was kept at 4°C as a non-sorted control, until further use. The remainder was stained using FITC-conjugated F(ab’)2-Goat anti-human IgA (Invitrogen; 1/1000) in PBS + 0.1% BSA (Fraction V; Roth) for 20 minutes at 4°C, washed once and resuspended in 100 μl PBS. All bacteria contained in the sample were subsequently sorted into FITC-positive and FITC-negative fractions, based on the FSC-A and anti-IgA-FITC parameters (**Figure 2A**), using a MA900 cell sorter (Sony) at the Flow Cytometry Core Facility Tübingen.

Nasal sIgA was obtained from 1 ml nasal swab eluates by centrifugation (1 minute, 10 000 xg) and supernatant was sterile-filtered through a 0.4 μm pore (Merck). We determined sIgA concentrations by ELISA (Human IgA ELISA kit; Thermo Fisher), replacing the provided monomeric IgA standards with sIgA ELISA standards (Abnova). All samples were stored at −20°C until use.

### DNA extraction, amplification and sequencing

We extracted DNA from the FITC-positive, FITC-negative and non-sorted fractions immediately after cell sorting using the QIAmp DNA Microbiome kit (Qiagen) and a FastPrep-24 Classic homogenizer (MP Biomedicals) for mechanical lysis of the bacterial cells, according to the manufacturers’ specifications. DNA was eluted in 50 μl of the supplied elution buffer, dried using a miVac centrifugal vacuum concentrator (SP Genevac), resuspended in 20 μl nuclease-free water (Ambion), and stored at −20°C until further use.

DNA amplification, amplicon library pooling and 16S rRNA sequencing were performed by the NGS Competence Center Tübingen (NCCT). Specifically, the 16S V1-V3 regions were amplified according to Escapa *et al*. (28), using primers 518F and 27R (**Table S2**). The pooled amplicon library was sequenced on the Illumina MiSeq platform using MiSeq Reagent Kit v3 (Illumina).

### Data processing

Demultiplexed reads were checked for primer presence using Cutadapt (v1.18) (39). We then used the DADA2 (v1.22.0) pipeline (40) in R (v4.1.3) (41) for raw reads quality filtering and trimming, error rate learning, sample inference, pair concatenation, ASV calling and chimera removal. The default parameters were used throughout, except for minParentAbundance=15 and minFoldParentOverAbundance=4 in the chimera removal step. The exported ASV table was imported in QIIME2 (v2021.11) (42) as a biom-formatted feature table. Taxonomic assignment was performed using a Naive-Bayes classifier trained as described previously using the eHOMD database (v15.1) (3, 28).

To calculate core diversity metrics, alpha-rarefaction curves were conducted in QIIME2. Then, using a sequence depth of 1100 reads (for analysis of the non-sorted samples only) or 800 reads (for all other analyses), matrices were computed using the Phyloseq (v1.38) (43) and Microbial (v0.0.20) (44) packages in R. Beta diversity metrics were demonstrated using weighted Unifrac distance and Principal Coordinate Analysis (PCoA) in QIIME2. IgA probability ratios were calculated using the IgAScores package in R (13). CST classification was performed using a custom Python script (deposited on GitHub: https://github.com/AhmedElsherbini/CST_picker), according to the original CST definitions (4).

### Bacterial strains and growth conditions

*S. aureus, S. epidermidis* and *S. lugdunensis* isolates (**Table S1**) were grown under aerobic conditions overnight at 37°C with agitation in 5 ml tryptone soy broth (TSB; Oxoid). *C. accolens* and *C. simulans* isolates **(Table S1**) were grown under anaerobic conditions for 42h at 37°C with agitation in brain-hearth infusion broth (BHI; Roth) with 0.4% Tween-80 (Sigma). *C. acnes* (**Table S1**) was grown under anaerobic conditions for 42h at 37°C on basic medium-blood agar (BM-blood; 10 g/l soy peptone A3SC (Organo Technie), 5 g/l yeast extract (Ohly), 5 g/l NaCl (Merck), 1 g/l K2HPO4-trihydrate (Thermo Fisher) 1 g/l D-glucose-monohydrate (Sigma-Aldrich), 5% defibrinated sheep blood (Oxoid), 1.5% agar (BD)), from which bacterial cells were collected prior to experiments. To simulate nutrient-limited conditions, *S. aureus* and *C. simulans* were grown as described above in 10% Todd-Hewitt broth (THB; Oxoid).

### Bacterial staining and flow cytometry

Bacteria were collected from triplicate cultures by centrifugation (1 minute, 10 000 xg) and resuspended at OD_600_ = 0.4 in PBS with 0.1% BSA. In case of *C. acnes*, bacterial cells were collected from agar plate using a sterile loop, resuspended in PBS with 0.1% BSA, and diluted to OD_600_ = 0.4. Bacteria were mixed 1:1 with sterile-filtered human nasal eluates diluted in PBS + 0.1% BSA to a final concentration of 0.3 μg/ml (for *S. aureus)* or 10 μg/ml sIgA (for all other species) and incubated for 30 min at 4°C. The samples were subsequently washed in PBS + 0.1% BSA, stained using FITC-conjugated F(ab’)2-Goat anti-human IgA (Invitrogen; 1/1000) in PBS + 0.1% BSA for 20 minutes, washed again and fixed with 1% formaldehyde (Sigma) in PBS. After 15 minutes, formaldehyde was washed off and the samples were resuspended in PBS. Per sample, 10 000 events were acquired on an LSRFortessa X-20 flow cytometer (BD Biosciences) and analyzed using FlowJo 10 software (BD Biosciences). The acquired geomean fluorescence intensities (FI) were normalized to the matching unstained control to facilitate comparison between the different species.

### Construction of *S. aureus* USA300 JE2 SpA-AA

The nucleotide sequence of the *S. aureus* USA300 FPR3757 *spa* gene (GenBank locus tag SAUSA300_RS00585) was used as a reference to create *spa-AA*, in which the D70A, D71A, D131A, D132A, D189A, D190A, D247A, D248A, D305A and D306A mutations were introduced (25). Additionally, we used alternative codons for residues 56 to 63 to introduce a unique primer annealing site for screening purposes. As the 293 bp upstream region of the *spa* start codon until the *spa* stop codon was too high in complexity due to highly repetitive regions, we had the region synthesized as two HiFi gBlocks (IDT), separated at the unique HindIII restriction site in the SpA A-domain. This reduced the template complexity of the first half sufficiently to synthesize it. For the second half, we introduced an additional 72 silent nucleotide substitutions to reduce the template complexity sufficiently to enable synthesis. Only frequently used codons (> 0.5%) were introduced, as listed in the Kazusa codon use database for *S. aureus* USA300 (45). Sequences for all gBlocks and primers are listed in **Table S2**.

Both gBlocks were PCR amplified using primers 1 and 2 (Biomers) for gBlock 1 and primers 3 and 4 for gBlock 2, digested at their unique HindIII (Thermo Fisher) restriction sites and ligated with T4 DNA ligase (Thermo Fisher), according to the manufacturers’ instructions. The ligated gBlock was subsequently digested and cloned into pIMAY (46) between the KpnI and SacI (Thermo Fisher) restriction sites, according to the manufacturers’ instructions, to create pIMAY-SpA-AA. This plasmid was transferred into *E. coli* IM08B (47) (**Table S1**) by heat shock procedure for plasmid amplification. Subsequently, *S. aureus* USA300 JE2 (**Table S1**) was transformed with pIMAY-SpA-AA and allelic exchange was performed as described by Monk *et al*. (46, 47). Successful exchange of the wildtype *spa* allele with the variant *spa-AA* allele was verified by PCR using primers 5 and 6, and Sanger sequencing (Eurofins) of the locus using primers 5, 7, 8 and 9.

### Statistical analyses

Normality of the data was tested by Shapiro-Wilk test. Statistical differences between two groups were tested by Mann-Whitney test, paired t-test or unpaired t-test. Differences in alpha diversity of multiple groups were tested by Kruskal-Wallis test with Dunn’s test for multiple comparisons. Correlations were analyzed by linear or non-linear regression analysis. IgA probability ratios were tested by Wilcoxon signed-rank test against a hypothetical value of 0. Significant differences are indicated by their exact p-values. All statistical analyses were performed using Graphpad Prism (v9.3.1) or the R stats package (41).

## Acknowledgements

The authors thank Esther Lehmann for methodological advice and support, Bernhard Krismer for critical feedback on the manuscript, the Flow Cytometry Core Facility (FCF-Berg, Universitätsklinikum Tübingen) for bacterial cell sorting, NGS Competence Center Tübingen (NCCT, Universitätsklinikum Tübingen) for 16S rRNA sequencing and methodological advice, and the High Performance and Cloud Computing Group (Zentrum für Datenverarbeitung, University of Tübingen) for computational resources.

This work received support from the Dutch Research Council (Rubicon grant 45219208 to R.v.D.), the European Molecular Biology Organization (EMBO fellowship ALTF 757-2019 to R.v.D.), and infrastructural support from the German Research Foundation (Cluster of Excellence ‘Controlling Microbes to Fight Infections’ EXC 2124 and bwForCluster BinAC INST 37/935-1 FUGG).

## Author contributions

Conceptualization and funding acquisition: R.v.D. and A.P.; formal analysis and visualization: R.v.D. and A.M.A.E.; investigation: R.v.D., M.H., S.A., and R.S.; methodology: R.v.D.; software: A.M.A.E.; supervision: A.P.; writing – original draft preparation: R.v.D.; writing – review & editing: R.v.D., A.M.A.E. and A.P.

## Supplementary figures

**Figure S1.**
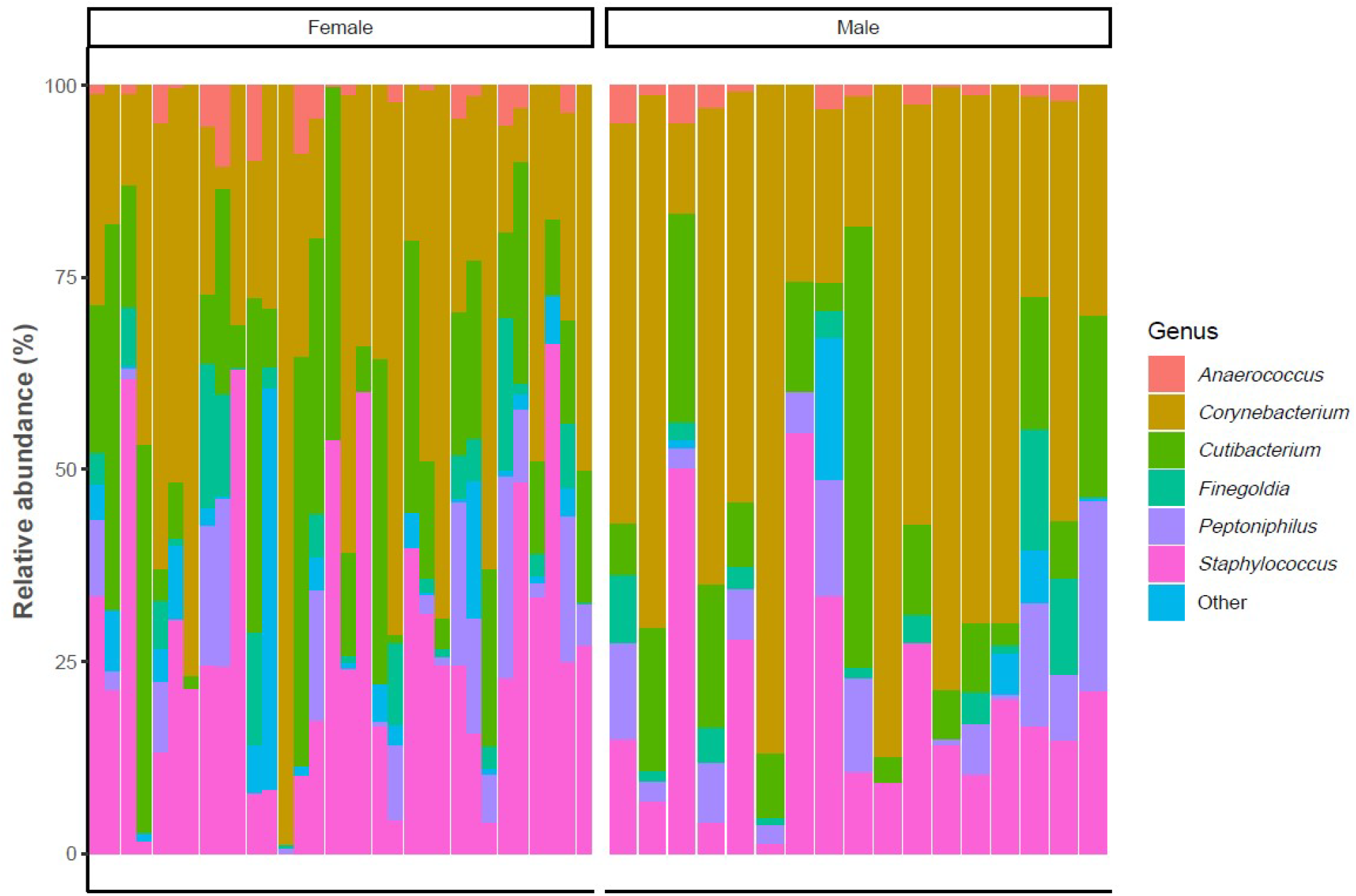
Compositional analysis by gender. Relative abundance of the five most abundant genera, stratified by gender.

**Figure S2.**
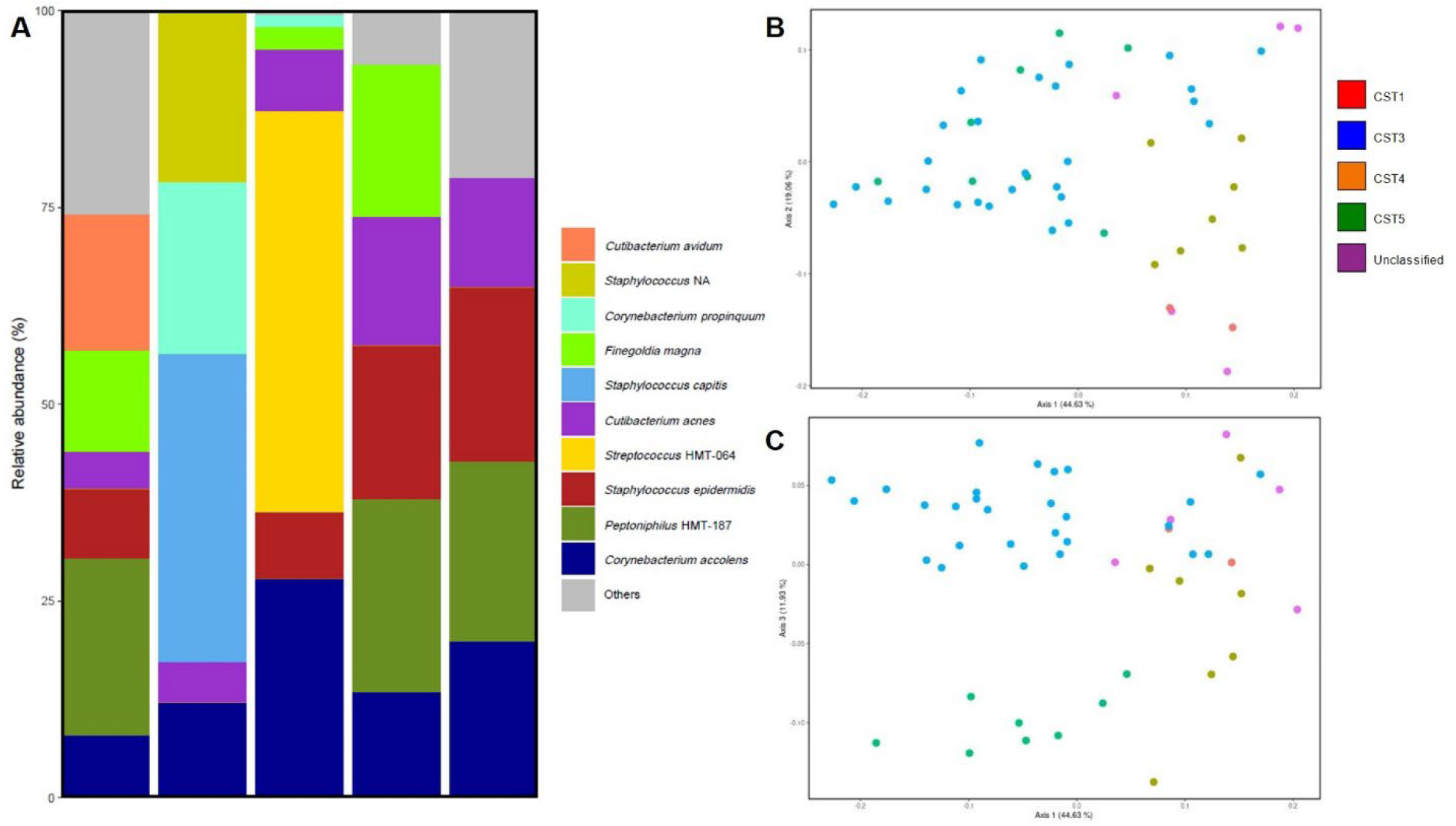
Analysis of the CST-unclassified microbiome samples. (**A**) Relative abundances of the most prevalent species of all CST-unclassified samples. *NA:* no species-level taxonomic classification could be determined. (**B-C**) Principal coordinates analysis of the beta diversity, expressed as the weighted Unifrac distance of (**B**) PCoA 1 (44.63%) vs. PCoA 2 (19.06%) and (**C**) PCoA 1 (44.63%) vs. PCoA 3 (11.93%).

**Figure S3.**
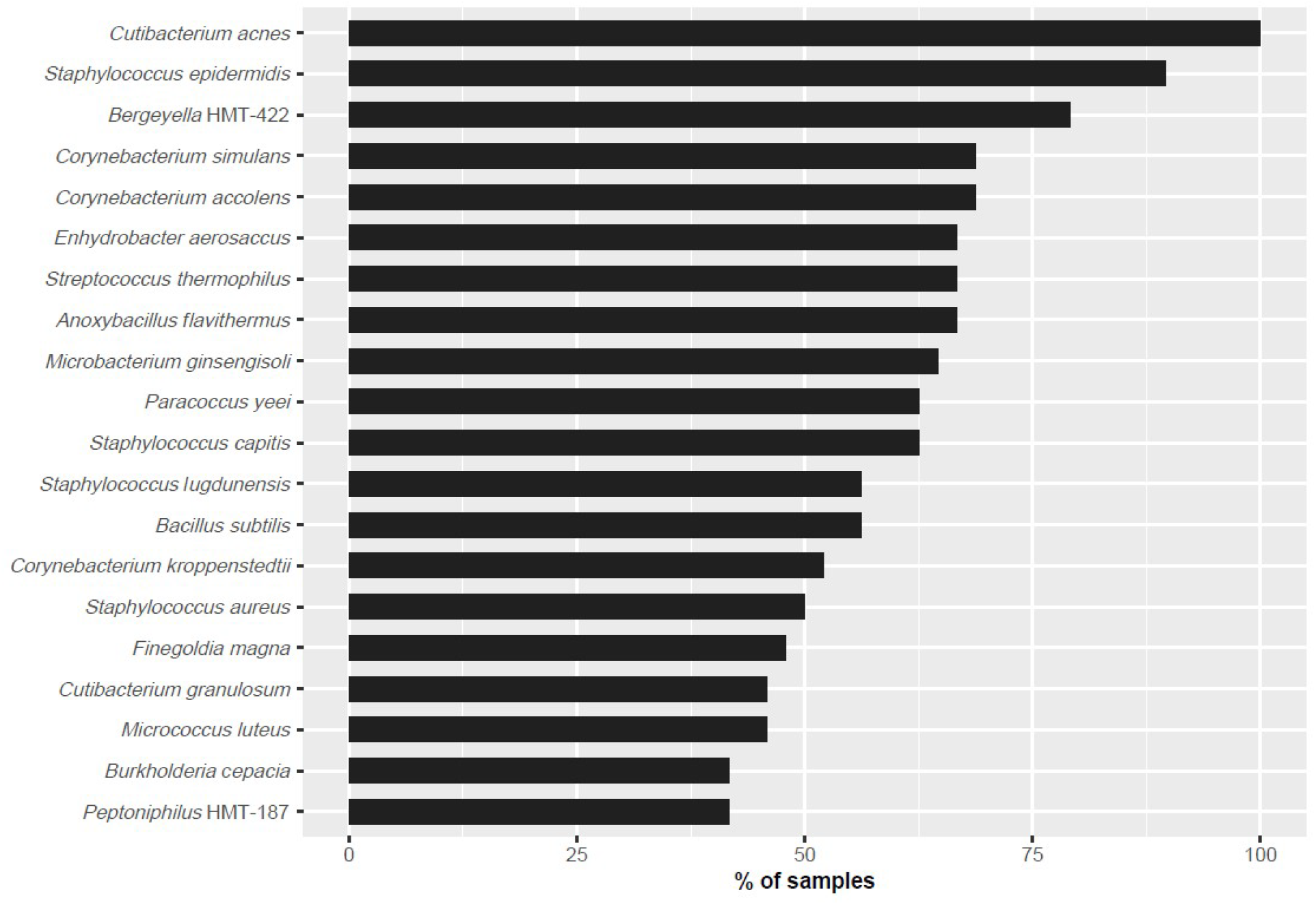
Top-20 species with the highest frequency of IgA scores. The frequency indicates in which percentage of the samples an IgA score for a given species could be calculated.

**Figure S4.**
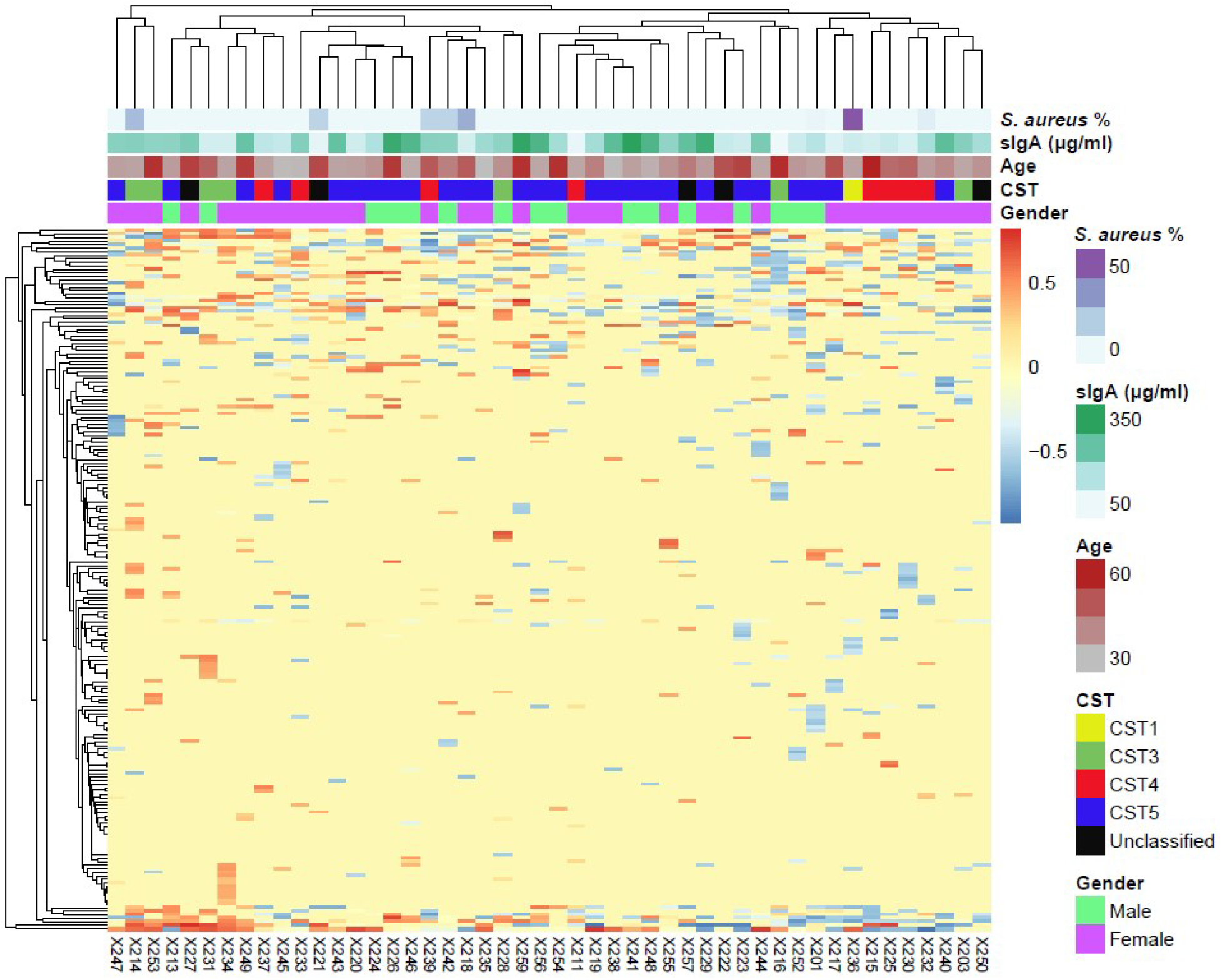
Hierarchical cluster analysis of the sIgA targeting. The study participants (columns) and all nasal bacterial species (rows) were clustered based on the IgA scores. Related to **Figure 4**.

## Supplementary file

**Supplementary file: IgA scores on genus and species level.**

## Supplementary tables

**Table S1.** Bacterial strains used in this study.

| Species | Strain | Source |
| --- | --- | --- |
| *Corynebacterium accolens* | 63VAs_B8 | Human nasal isolate (48) |
| *Corynebacterium simulans* | 50VAs_B5 | Human nasal isolate (48) |
| *Cutibacterium acnes* | Mü58 | Human nasal isolate (48) |
| *Escherichia coli* | IM08B | Cloning host for plasmid transfer to *S. aureus* (47) |
| *Staphylococcus aureus* | 35-1 | Human nasal isolate from in-house collection |
| *Staphylococcus aureus* | JE2 | Human abscess isolate, USA300 clone (49) |
| *Staphylococcus aureus* | JE2 SpA-AA | This study |
| *Staphylococcus epidermidis* | IVK83 | Human nasal isolate (5) |
| *Staphylococcus lugdunensis* | IVK28 | Human nasal isolate (50) |

**Table S2.**
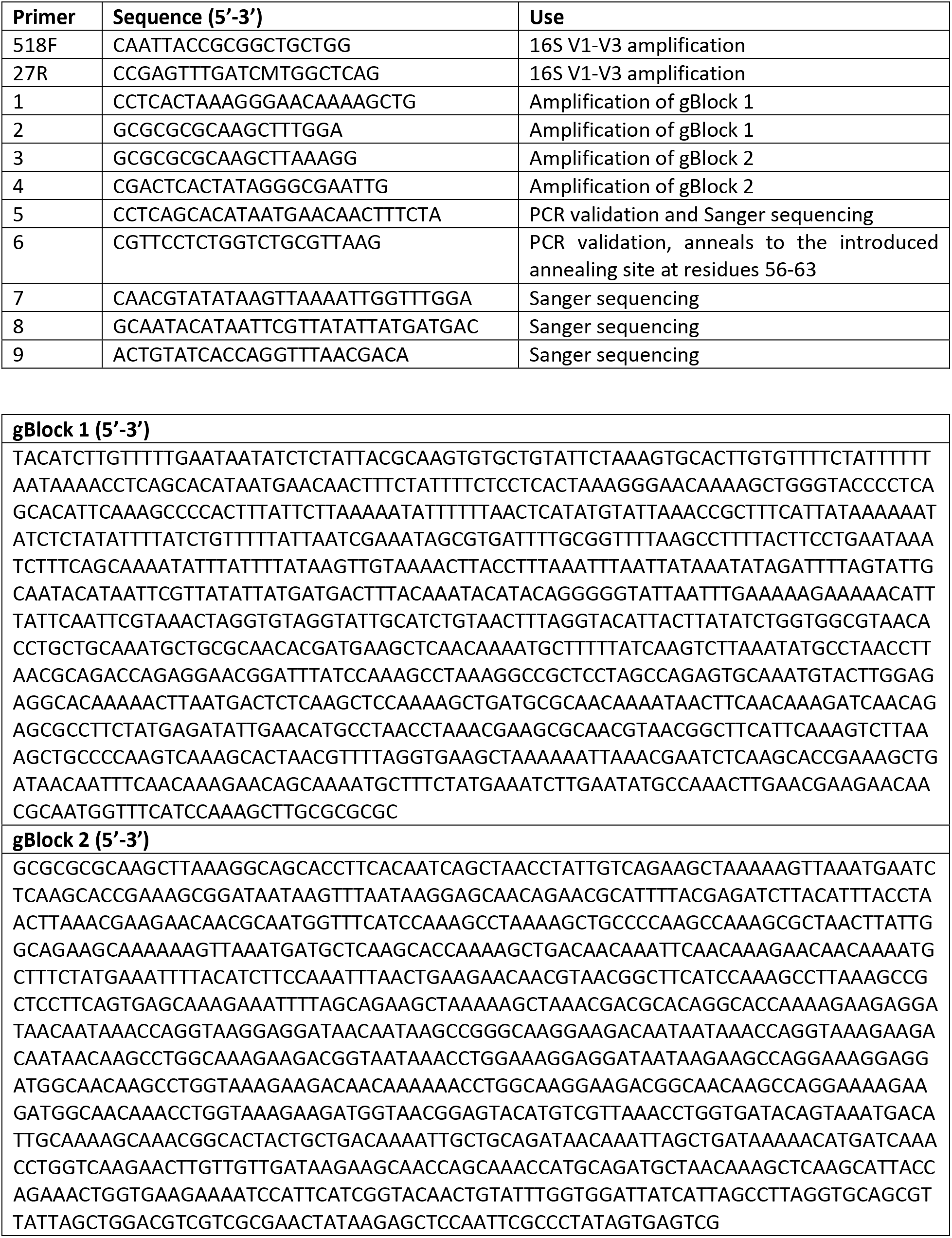
Primers and gBlocks used in this study.

